# Loss of Cdc42 causes abnormal optic cup morphogenesis and microphthalmia in mouse

**DOI:** 10.1101/2024.10.20.619331

**Authors:** Katrina S. Hofstetter, Paula M. Haas, Jonathon P. Kuntz, Yi Zheng, Sabine Fuhrmann

**Author notes:** <u>Corresponding author:</u> Sabine Fuhrmann, Dept. of Ophthalmology and Visual Sciences, Vanderbilt University Medical Center, 1161 21^st^ Ave S, AA7100 MCN (in VUIIS), Vanderbilt Eye Institute Research Laboratories, Nashville, TN 37232, Phone: 615/936-0621. Division of Infectious Diseases, Department of Medicine, Emory University, Atlanta, Georgia, USA. Department of Life and Environmental Sciences, University of California Merced, Merced, CA, USA.

## Abstract

Congenital ocular malformations originate from defective morphogenesis during early eye development and cause 25% of childhood blindness. Formation of the eye is a multi-step, dynamic process; it involves evagination of the optic vesicle, followed by distal and ventral invagination, leading to the formation of a two-layered optic cup with a transient optic fissure. These tissue folding events require extensive changes in cell shape and tissue growth mediated by cytoskeleton mechanics and intercellular adhesion. We hypothesized that the Rho GTPase Cdc42 may be an essential, convergent effector downstream of key regulatory factors required for ocular morphogenesis. CDC42 controls actin remodeling, apicobasal polarity, and junction assembly. Here we identify a novel essential function for Cdc42 during eye morphogenesis in mouse; in *Cdc42* mutant eyes expansion of the ventral optic cup is arrested, resulting in microphthalmia and a wide coloboma. Our analyses show that Cdc42 is required for expression of the polarity effector proteins PRKCZ and PARD6, intercellular junction protein tight junction protein 1, β-catenin, actin cytoskeleton F-actin, and contractile protein phospho myosin light chain 2. Expression of RPE fate determinants OTX2 and MITF, and formation of the RPE layer are severely affected in the temporal domain of the proximal optic cup. EdU incorporation is significantly downregulated. In addition, mitotic retinal progenitor cells mis-localized deeper, basal regions, likely contributing to decreased proliferation. We propose that morphogenesis of the ventral optic cup requires Cdc42 function for coordinated optic cup expansion and establishment of subretinal space, tissue tension, and differentiation of the ventral RPE layer.

## Introduction

Eye morphogenesis is a highly dynamic process; first optic vesicles evaginate from the ventral forebrain, comprised of specified neuroepithelial progenitor cells determined to become optic stalk, neural retina, and retinal pigment epithelium (RPE; Supplementary Figure 1A). Subsequently, the distal and ventral optic vesicle and the overlying surface ectoderm undergo invagination, resulting in a two-folded optic cup, a lens vesicle, and the ventral optic fissure. The optic fissure then fuses resulting in a continuous optic cup. Disruption of any of these processes leads to congenital ocular malformations producing ∼25% of childhood blindness: microphthalmia (small eye), anophthalmia (absent eye), and coloboma (optic fissure closure defect in the ventral optic cup), collectively hereafter MAC (Clementi et al., 1992; Graw, 2019; Morrison et al., 2002). These morphogenetic processes require dynamic changes in cell and tissue shape through regulation of apicobasal polarity, cell adhesion, cytoskeleton dynamics, growth and tissue-tissue interaction, as well as acquisition of regionalized cell fate for establishment of the retina, RPE and optic stalk (Cardozo et al., 2023; Casey et al., 2021; Chan et al., 2020; Fuhrmann, 2010; Hosseini and Taber, 2018; Miesfeld and Brown, 2019). Consistent with a prominent role of actin cytoskeleton dynamics during optic fissure closure, coloboma is found in patients of Baraitser-Winter Syndrome with mutations in the actin gamma cytoplasmic 1 gene (*ACTG1)* causing reduced incorporation into filamentous actin (F-actin) (Rainger et al., 2017). While regionalization of optic cup and optic vesicle are well studied (Diacou et al., 2022; Fuhrmann, 2010; Miesfeld and Brown, 2019; Viczian, 2014), the role of effectors regulating changes in cell and tissue morphology is not well understood.

The small Rho GTPase cell division cycle 42 (Cdc42) controls establishment, remodeling and maintenance of epithelia during development and homeostasis (Duquette and Lamarche-Vane, 2014; Rolo et al., 2018; Zhang et al., 2022). Cdc42 instructs apicobasal polarity and is critical for localization of cytoskeleton components, intercellular junctions, mitosis, and filopodia formation (Mack and Georgiou, 2014; Pichaud et al., 2019; Sit and Manser, 2011). Cdc42 activates protein kinase C (PKC) and par-6 family cell polarity regulator (Pard6) to form the apical polarity complex, regulating segregation of apical and basal polarity components, establishing junctional complexes, and it activates the effector Cdc42 binding protein kinase (MRCK) to promote apical actomyosin contractility. In mice, early disruption of *Cdc42* in telencephalic neuroepithelial cells results in polarity and differentiation defects, hyperplasia and holoprosencephaly (Cappello et al., 2006; Chen et al., 2006). During optic cup morphogenesis, Cdc42 is expressed and participates in regulating lens pit invagination, by inducing filopodia to couple lens vesicle and retina (Chauhan et al., 2009; Mitchell et al., 2007). During retinal neurogenesis, Cdc42 is essential for lamination and tissue integrity, likely by promoting assembly of adherens junction complexes (Heynen et al., 2013). In zebrafish *Cdc42* morphants, smaller eyes, mild lamination defects, absence of photoreceptor cilia and decreased survival is observed (Choi et al., 2015; Choi et al., 2013). Loss of *Cdc42* during early eye morphogenesis appears to lead to a delay in eye development, however, this has not been further investigated (Chen et al., 2006). Here, we examined the early role of Cdc42 in eye morphogenesis, by performing temporally controlled Cre-mediated ablation. Our observations reveal a novel role for Cdc42; it is essential for growth, proliferation, differentiation and optic fissure closure during optic cup morphogenesis.

## Methods

Animal procedures were reviewed and approved by the Institutional Animal Care and Use Committee at Vanderbilt University Medical Center. Mouse strains were maintained in the C57BL/6J background. To generate *Cdc42* mutant embryos, mice harboring conditional *Cdc42* and recombination reporter *RosaR26* (*Gt(ROSA)26Sor*^*tm1Sor*^, Jax # 3474) alleles were crossed with *Hes1*^*tm1(cre/ERT2)Lcm*^ (hereafter *Hes1*^*CreERT2*^) (Chen et al., 2006; Kopinke et al., 2011; Soriano, 1999). Genotyping was performed with established protocols and by Transnetyx (Cordova, TN) using Taqman with custom-designed probes. Noon of the day with an observed vaginal plug was counted E0.5. Pregnant dams received 0.05-0.15 mg/g tamoxifen (Sigma T5648) by oral gavage between E7.5 and E8.5. For analysis of proliferation, pregnant dams received one intraperitoneal EdU injection two hours before sacrificing (30 µg/g; Invitrogen E10187). We routinely confirm for absence of *Crb1*^*Rd8*^ in our mouse colony, particularly in the strains used for this study.

Mutant embryos with conditional deletion of *Cdc42* (hereafter *Cdc42*^*CKO*^) and control littermates were processed as previously published (Sun et al., 2020). For antigen retrieval, coronal or sagittal cryostat sections were treated with 1% Triton X-100. The following primary antibodies were used: β-catenin 1:3,000 (Sigma-Aldrich; Darmstadt, Germany; #C2206), β-galactosidase 1:4,000, Cappel; MP Biomedicals, Aurora, OH; #55976, phospho Jun terminal kinase (pJNK), 1:750, Promega; Madison, WI; #V7931, microphthalmia associated transcription factor (MITF), 1:800, Exalpha; Shirley, MA; # X1405M, phospho myosin light chain 2 (pMLC2), 1:80, Cell Signaling; Danvers; #3674, OTX2, 1:1,500, R&D Systems; Minneapolis, MN; #AF1979, PARD6, 1:300, Santa Cruz Biotechnology; Dallas, TX; #sc-67393, paired box 2 (PAX2), 1:800, BioLegend; San Diego, CA; #901001, paired box 6 (PAX6), 1:500, BioLegend; San Diego, CA; #901301, phospho histone H3.1 (pH3.1), 1:2,500, Sigma-Aldrich; Darmstadt, Germany; #H9908, protein kinase C zeta (PRKCZ), 1:500, Santa Cruz Biotechnology; Dallas, TX; #sc-216, visual system homeobox 2 (VSX2), 1:800, Exalpha; Shirley, MA; #X1180P, TJP1, 1:500, Invitrogen/ThermoFisher; Walham, MA; #61-7300. The following secondary antibodies were used: donkey anti-mouse Alexa Fluor®647, 1:800, Thermo Fisher Scientific; Walham, MA; #A31571, donkey anti-rabbit Alexa Fluor®488, 1:1,000, Jackson ImmunoResearch; West Grove, PA; #711-545-152, donkey anti-rabbit Alexa Fluor®647, 1:500, Jackson ImmunoResearch; West Grove, PA; #711-605-152, donkey anti-rat Alexa Fluor®488, 1:1,000, Jackson ImmunoResearch; West Grove, PA; #712-545-150, donkey anti-goat Alexa Fluor®568, 1:1,000, Thermo Fisher Scientific; Walham, MA; #A11057, donkey anti-goat Alexa Fluor®647, 1:1,000, Thermo Fisher Scientific; Walham, MA; #A21447, donkey anti-goat Rhodamine Red® RED, 1:800, Jackson ImmunoResearch; West Grove, PA; #705-295-147. Filamentous actin (F-actin) was detected using Phalloidin (1:75; Thermo Fisher Scientific A12379). ApopTag Fluorescein In Situ Apoptosis Detection Kit (EMD Millipore S7110) was used to detect apoptotic cells. For EdU detection, the Click-iT® EdU Imaging Kit (Thermo Fisher Scientific C10637) was utilized. Cryostat sections were counter-labeled with DAPI and mounted in Prolong Gold Antifade. No developmental defects were observed in conditional heterozygous female (hereafter *Cdc42*^*CHET*^) or *Cdc42*^*FL/FL*^ embryos without *Cre* (hereafter Con). Unless otherwise indicated, at least 3 embryos from a minimum of 2 individual litters were analyzed per genotype, time point, and marker.

Images were captured using an Olympus SZX12 stereomicroscope, equipped with U-CMAD3 camera, and Olympus BX51 epifluorescence system (XM10 camera). We used Olympus FV100 or ZEISS LSM 880 systems for confocal imaging. Images were processed using ImageJ (NIH, v.2.9) and Adobe Photoshop software (version 25.9.0). In images showing sagittal orientation of embryo heads, temporal is located on the right.

### Quantification of eye size, subretinal space and shortening, cellular organization, cell shape and cell alignment

For quantification of eye size, the circumferential area encompassing the eye along the basal border of the RPE was traced (average of 3 measurements per eye) on bright field images captured with a SZX12 stereomicroscope at highest magnification (90x magnification, 1x objective). The area in the region of interest (ROI) was calculated using ImageJ. In *Cdc42*^*CKO*^ embryos, the ventral colobomatous gap was traced by extension from the pigmented fissure edges and below or along the ventral lens vesicle boundary (see example in Supplementary Figure 1H inset). The subretinal space was defined as the space between the apical boundaries of retina and RPE. Length and shortening of subretinal space were measured on epifluorescence images using ImageJ (average of 3 measurements per eye; Supplementary Figure S2F).

Cellular disorganization was outlined by visual observation in 2 proximal phalloidin- and beta-catenin-labeled sections per embryo of 4 control and 4 *Cdc42*^*CKO*^ (see example in Supplementary Figure 2I) and measured using ImageJ. We quantified cell shape by measuring length and width of 21-25 cells per embryo adjacent to the patterning defects in the presumptive RPE layer of the ventral optic cup (temporal side, proximal level). Examples are shown in Supplementary Figure 2K, by magnification of the boxed region in the ventral temporal optic cup (see 2I). To investigate cell alignment, we measured in 21-25 cells per embryo the angle between the length of each cell and the basal boundary of the optic cup (set as 0 degrees; ImageJ; Supplementary Figure 2K; 4 control and 4 *Cdc42*^*CKO*^). Any deviation from a 90 degree-angle, either higher or lower, was expressed as an according value between 0 and 90. For example, an angle of 130 degree was converted to 50 degrees.

### Quantification of cell death, proliferation and cell position

We quantified TUNEL- and EdU-labeled cells in 2-3 (TUNEL) and 2-5 (EdU) coronal sections per embryo midway through the optic cup as a percentage of total DAPI-labeled nuclei. For EdU analysis, 7 control, 6 *Cdc42*^*HET*^ and 5 *Cdc42*^*CKO*^ embryos were analyzed. For TUNEL analysis, 6 control and 6 *Cdc42*^*CKO*^ embryos were examined. The same sections were co-labeled for detection of OTX2 protein, and images were captured using the BX51 epifluorescence system.

Samples lacking ventral RPE were omitted from the analysis for ventral RPE. For apical distance measurements in sagittal sections, the shortest distance between pH3.1-labeled nuclei of retinal progenitors and apical boundary of the retina was measured on epifluorescence images using ImageJ. Analyzed were for E10.5, 2-4 sections per embryo (5 control and 3 *Cdc42*^*CKO*^ embryos), for E11.5, 2-3 section per embryo (4 control, 3 *Cdc42*^*CKO*^ embryos), and for E12.5, 2-4 sections per embryo (4 control, 4 *Cdc42*^*CKO*^ embryos).

Prism 10 (Graphpad) was used for statistical analysis. A p-value below 0.05 was considered as statistically significant. Unless otherwise specified, measurement values were expressed as mean ± standard deviation.

## Results

### Conditional *Cdc42* inactivation at the eye field stage (E7.5 - E8.0) does not interfere with optic vesicle morphogenesis and initial invagination

We performed temporally controlled, conditional inactivation by *Hes1*^*CreERT2*^ to determine when exactly Cdc42 is required during ocular morphogenesis. Analysis of *RosaR26* reporter expression confirmed *Hes1*^*CreERT2*^-mediated recombination in the retina, with mosaic activity in RPE, lens and in some extraocular mesenchyme cells (Supplementary Figure S1B-D; (Yun et al., 2009). Thus, any loss of CDC42 expression in lens and extraocular mesenchyme may contribute to ocular developmental abnormalities. We tried several CDC42 antibodies unsuccessfully, thus it is unclear whether CDC42 is absent in cells expressing *RosaR26* reporter in an identical spatial and temporal pattern. To investigate a potential role in optic vesicle formation and invagination, we first induced recombination around the eye field stage (E7.5 - E8.0) and harvested embryos at the time points E10.5 and E11.0. In *Cdc42*^*CKO*^ embryos, we did not observe severe defects in early eye morphogenesis, confirmed by localization of F-actin (Figure 1B, D; embryos harvested at E11.0). Early invagination of the optic vesicle and lens ectoderm did occur in mutant eyes, indicating that Cdc42 is not required for initiating morphogenesis of optic and lens vesicles.

**Figure 1:**
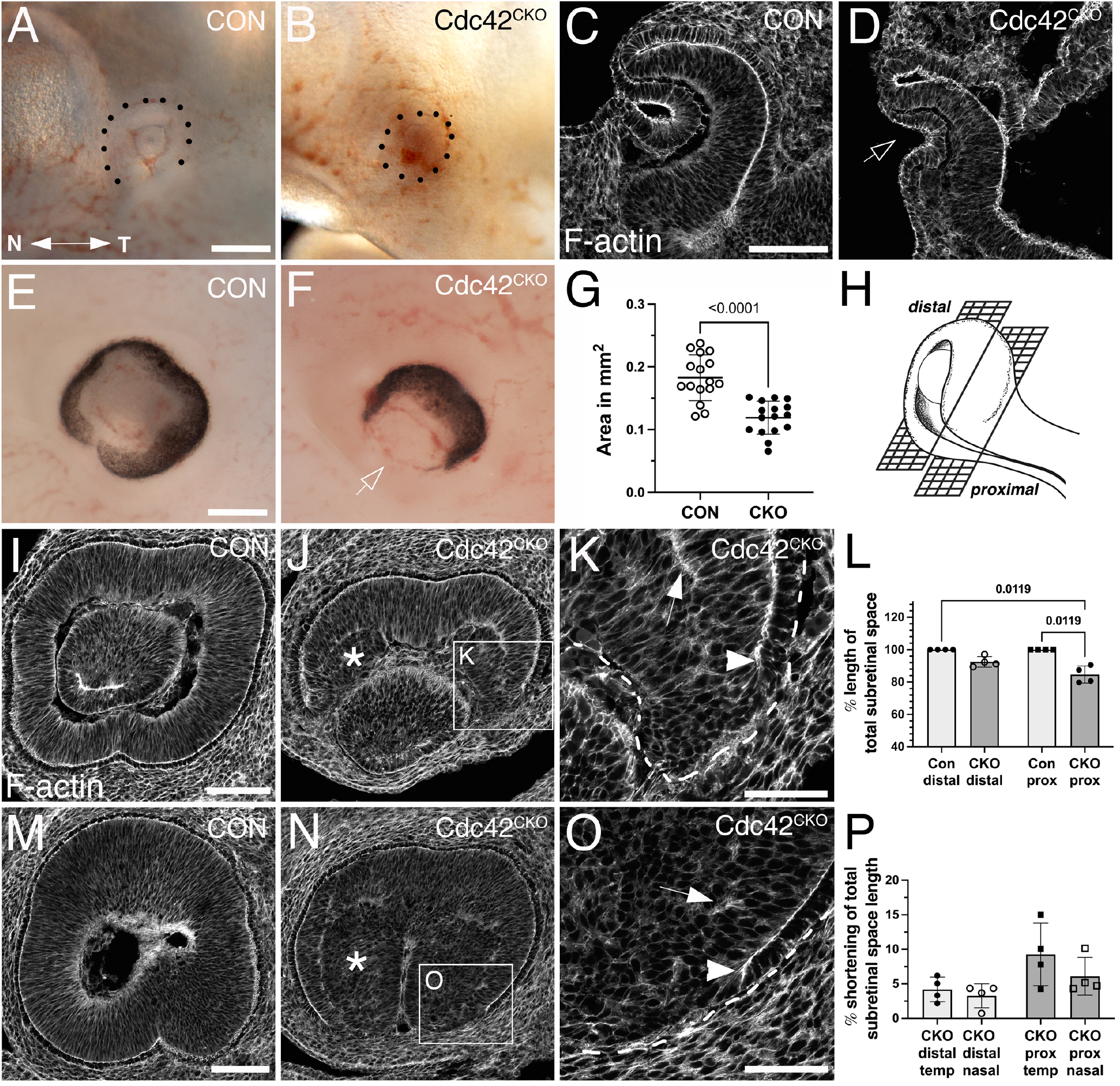
Effects of Cdc42 disruption between E7.5 and E8.5 on eye morphogenesis. A-D) *Cdc42* ablation between E7.5-E8.0 does not interfere with eye morphogenesis until formation of the optic cup starts, here shown in embryos harvested at E11.0. (A) Control (*Cdc42*^*FL/FL*^, 33 somites). (B) *Cdc42*^*CKO*^ (31 somites). Dotted lines mark the circumference of the eye. Temporal right, nasal left. (C, D) Phalloidin labeling in control (C, *Cdc42*^*FL/FL*^, 33 somites) and *Cdc42*^*CKO*^ optic cups (D, 31 somites) in coronal orientation shows invagination, including the overlying lens ectoderm (D, arrow) (4 and 7 embryos analyzed for control and *Cdc42*^*CKO*^, respectively. (E-P) *Cdc42* disruption at E8.5 interferes with morphogenesis of the ventral optic cup in E12.5 embryos. Temporal right, nasal left. (E, F) Control with normal optic cup (E, *Cdc42*^*FL/FL*^) and *Cdc42*^*CKO*^ showing a wide coloboma (F, arrow). G) Quantification of the circumferential area of left and right eyes at E12.5 shows a significant decrease by 35% in *Cdc42*^*CKO*^ (data points represent 16 eyes from 8 embryos per genotype). Unpaired T-test. (H) Cartoon showing distal and proximal levels of the optic cup in sagittal orientation. In sagittal views, nasal and temporal orientation are left and right, respectively. (Created in Biorender.com) (I-K) Phalloidin labeling of distal sagittal sections reveals loss of ventral tissue in *Cdc42*^*CKO*^ (J), compared to control (I, *Cdc42*^*FL/FL*^). Pseudostratified radial organization of retinal progenitors is disrupted in the inner optic cup (J, asterisk). (K) Higher magnification of inset in (J) reveals premature stop of apical boundaries in the edges of *Cdc42*^*CKO*^ optic cups (arrowhead) and F-actin mis-localization (arrow). (L) Quantification of apical boundary length in the subretinal space, shown as percentage of total subretinal space (= fully extended to optic cup margins). Kruskal-Wallis test, n=4 embryos per genotype. M-O) Phalloidin labeling of proximal sagittal orientation shows abnormal early termination of the apical boundaries in *Cdc42*^*CKO*^ (O, arrowhead), compared to control (M, *Cdc42*^*FL/FL*^). Pseudostratified organization of retinal progenitors is disrupted in the inner optic cup (N, asterisk) and F-actin is mis-localized (O, arrow). P) Quantification of apical boundary shortening in the subretinal space at distal and proximal level in *Cdc42*^*CKO*^, shown as percentage of total subretinal space (= fully extended to optic cup margins). One-way ANOVA with Tukey’s posthoc analysis, n=4 embryos per genotype. Scale bars A, E: 0.2 mm; C, I, M: 100 µm; K, O: 50 µm.

However, *Cdc42*^*CKO*^ embryos showed a general developmental delay when harvested beyond E10.5 (Supplementary Figure S1F; 4 of 7 *Cdc42*^*CKO*^ embryos, = 57%) (Park et al., 2008).

### Conditional *Cdc42* inactivation at E8.5 results in defective morphogenesis in the ventral optic cup

To investigate a requirement for *Cdc42* during subsequent optic cup morphogenesis, we administered tamoxifen later, at E8.5, and analyzed embryos at E12.5 when optic cup invagination and optic fissure closure is completed. *Cdc42*^*CKO*^ embryos appeared to develop normally but exhibited a severe tissue loss in the ventral optic cup, resulting in a wide coloboma (Figure 1F; Supplementary Figure S1H). Quantification of the eye circumference (see Methods for detailed description) confirmed microphthalmia in *Cdc42*^*CKO*^ embryos with a significant reduction by 35% (Figure 1G, Supplementary Figure 1H; 16 eyes from 8 embryos per genotype). Consistent with decreased eye size, we observed in Dapi-labeled consecutive sections a severe loss of ventral tissue in the distal optic cup and a dramatic reduction of vitreal space proximally (compare Supplementary Figure 2A-D with E-H). Retinal progenitor cells normally organize as a pseudostratified epithelial layer (Figure 1I, M, Supplementary Figure S2A-D). Nuclear labeling of *Cdc42*^*CKO*^ eyes showed disorganization of progenitor cells in inner optic cup regions (Supplementary Figure S2E-H), confirmed by phalloidin labeling (Figure 1J, K, N, O). In addition, irregular F-actin localization was observed (Figure 1K, O). We traced disorganized regions in the proximal optic cup of phalloidin- and β-catenin-labeled sections and measured their percentage of total eye area (Supplementary Figure 2I, J). Quantification showed a significant occurrence of disorganized regions in *Cdc42*^*CKO*^ optic cups.

Phalloidin labeling of distal and proximal levels of the optic cup (Figure 1H) showed loss of F-actin and apical boundaries close to the optic fissure (Figure 1J, K, N, O). The subretinal space, defined as the space between the apical boundaries of retina and RPE layers, was severely shortened and did not extend into the optic fissure margins in *Cdc42*^*CKO*^, compared to controls (Figure 1I, L, M). Thus, a defined RPE layer is discontinued, and this RPE loss occurred mostly in the proximal optic cup and more frequently in the temporal domain (Figure 1P). In summary, Cdc42 is required for expansion of ventral tissue and subretinal space that is necessary for apposition of the optic cup margins to establish a distinct optic fissure and achieve subsequent tissue fusion.

Absence of RPE correlates with loss of apical polarity in the ventral optic cup of *Cdc42*^*CKO*^In addition to establishing apicobasal polarity, proper regionalization of optic cup domains is an essential prerequisite for eye morphogenesis to proceed normally. The shortening of subretinal space in the ventral *Cdc42*^*CKO*^ optic cup suggested effects on the differentiation of RPE cells that normally extend into the optic fissure margins. Double-labeling with the RPE key regulatory transcription factor OTX2 and phalloidin (Figure 2A-D) or with polarity proteins PARD6 and pMLC2 (Figure 2E-H) confirmed that the loss of subretinal space is correlated with a reduction or complete loss of OTX2-labeled RPE precursor cells in the proximal temporal optic cup (Figure 2B, D, F, H). Furthermore, at higher magnification disorganized F-actin and loss of radial organization of retinal progenitor cells in *Cdc42*^*CKO*^ optic cups is detectable (Figure 2D). Co-labeling for the Cdc42 effector PRCZ and the retinal progenitor-specific protein VSX2 revealed that retinal differentiation did not expand into the ventral optic cup that is normally occupied by RPE precursors (Figure 2J, L). Another key regulatory transcription factor for RPE differentiation, MITF, was absent in the ventral optic cup of *Cdc42*^*CKO*^ (Figure 2N, P). The pan-ocular protein PAX6 appeared to be normally expressed throughout the ventral optic cup (Figure 2N). PAX6 was also present in the dorsal forebrain, thus, it is unclear whether and how the cells in the defective RPE layer are re-specified. PAX6 labeling further confirmed that subretinal space is discontinuous and that retinal progenitor nuclei appeared disorganized and rounder, in contrast to their pseudostratified arrangement in control (Figure 2M, N). Furthermore, the transcription factor PAX2 normally starts to be expressed in the proximal optic cup but was excluded from this region in *Cdc42*^*CKO*^ (Figure 2Q, R). PAX2 was still present in the optic stalk in *Cdc42*^*CKO*^ (not shown). Effects on localization of the tight junction protein TJP1 further confirmed polarity defects in the ventral subretinal space (Figure 2Q, R). Cdc42 forms complexes with cadherin proteins to regulate polarity, and the protein β-catenin is an essential structural component of adherens junctions coordinating catenin-cadherin complexes. It is normally expressed in apical boundaries of retina and RPE encompassing the subretinal space.

**Figure 2:**
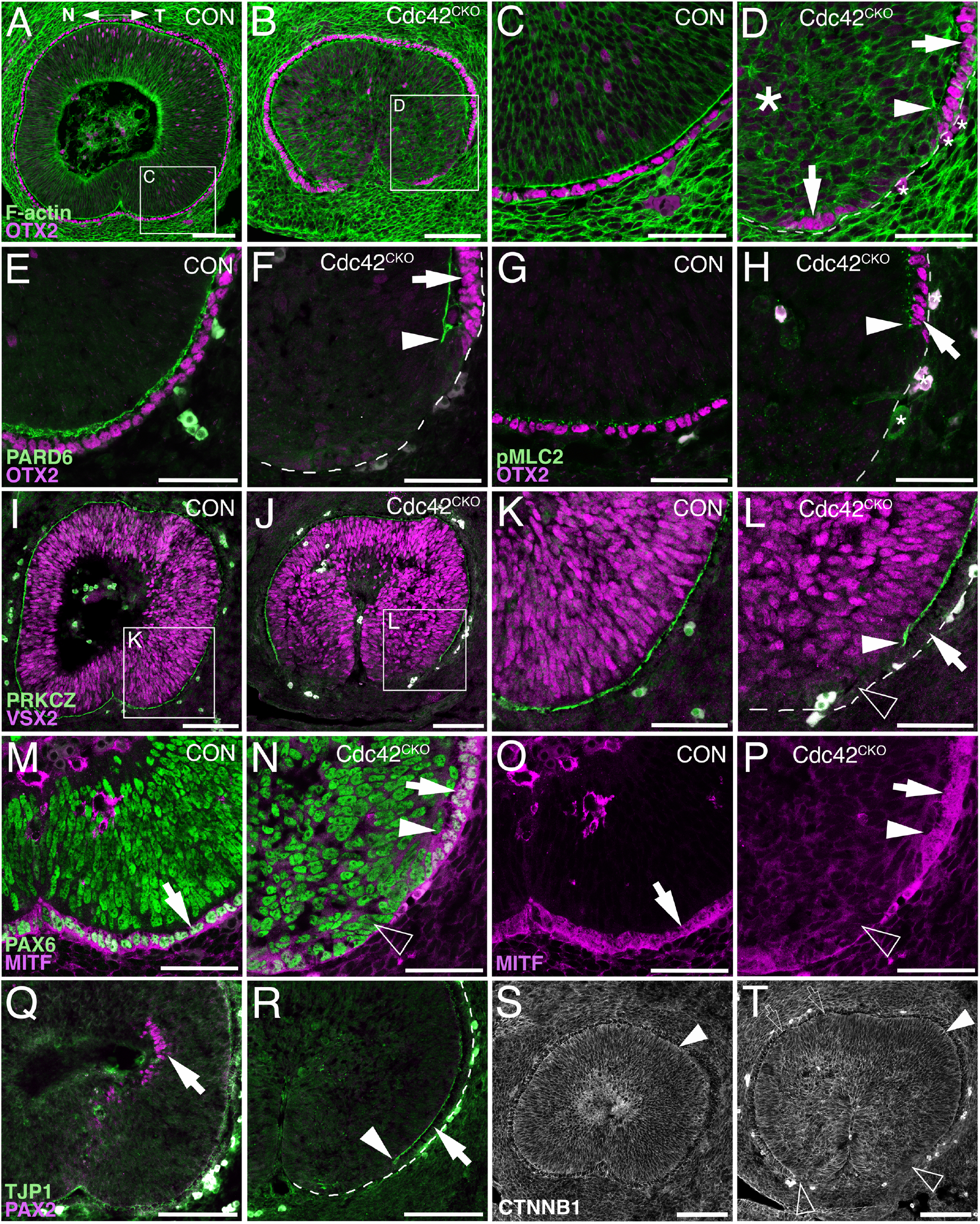
Defective polarity protein localization and loss of RPE tissue regionalization in the ventral *Cdc42*^*CKO*^ optic cup. A-P) Sagittal view of E12.5 proximal optic cup in controls (A, C, E, G, I, M, O, *Cdc42*^*FL/FL*^) and *Cdc42*^*CKO*^ (B, D, F, H, J, L, N, P). Nasal and temporal are oriented left and right, respectively. (A) In controls, OTX2 is expressed in the RPE layer (magenta) and F-actin is present as a distinct apical boundary in the subretinal space (green). Both are continuous in the ventral optic cup with completed fusion of the optic fissure. (B) In *Cdc42*^*CKO*^, the ventral optic cup has not fused, and OTX2 labeling is discontinuous on the temporal side (inset). (C, D) High magnification of insets from (A, B). In *Cdc42*^*CKO*^, the apical boundaries of retina and RPE encompassing the subretinal space are disrupted far up into the optic cup (D, arrowhead). The arrows point to OTX2-labeled RPE cells and small asterisks label cells with non-specific staining (D). Large asterisk on the left marks area of tissue disorganization (D). (E-H) Colocalization of OTX2 (magenta, arrows) and PARD6 (E-F, green) or pMLC2 (G-H, green) at high magnification in the temporal ventral optic cup. (F, H) In *Cdc42*^*CKO*^, the apical boundaries do not extend into the ventral optic cup (arrowheads), arrows point to OTX-labeled cells adjacent to the region of intact subretinal space. (I-L) Double labeling for VSX2 (magenta) and PRKCZ (green), and high magnification of insets from (I, J) are shown in (K, L). Shortening of the apical boundaries lining the subretinal space can be observed in *Cdc42*^*CKO*^ (J, L, arrowhead). VSX2 expression is not upregulated in ventrally located cells lacking the apical boundary in *Cdc42*^*CKO*^ (L, open arrowhead). (M-N) Co-labeling for PAX6 (green) and the RPE marker MITF (magenta) in the temporal ventral optic cup. (O-P) Single labeling for MITF shown in (M-N). In *Cdc42*^*CKO*^, PAX6 is expressed in cells occupying the region becoming normally the RPE layer (N, open arrowhead). Arrows points to MITF/PAX6-colabeled RPE cells (M-P), arrowhead marks the premature stop of the subretinal space (N, P). (Q-R) Temporal ventral optic cup showing double labeling for PAX2 (magenta) and TJP1 (green). In *Cdc42*^*CKO*^, PAX2-labeled cells are not detectable (R), compared to control (Q, arrow). (R) Arrow points to the RPE layer adjacent to intact subretinal space, arrowhead marks the end of the subretinal space. (S-T) Labeling for β-catenin in control (S) and *Cdc42*^*CKO*^ (T) embryos did not show obvious differences. Apical localization of β-catenin appeared maintained in *Cdc42*^*CKO*^ (filled arrowheads) and absent corresponding to absence of subretinal space (open arrowheads). Very small patches with cellular disorganization can be present in dorsal regions adjacent to the subretinal space, which do not result in defects of subretinal space formation (arrows). Scale bars A, B, I, J, O, P, S, T: 100 µm; C, D, E-H, K-N: 50 µm.

Besides loss of expression in the region normally occupied by subretinal space, *Cdc42*^*CKO*^ embryos did not show obvious differences in β-catenin distribution (Figure 2T). Apical localization of β-catenin appeared maintained in the dorsal optic cup of *Cdc42*^*CKO*^. Very small patches with cellular disorganization can be found in dorsal regions, which do not result in defects in subretinal space formation (Figure 2T). Thus, in contrast to the ventral optic cup, dorsal expression of polarity and adhesion markers appeared unaffected in Cdc42^CKO^ (Figure 1J, N, 2B, J, R, T).

Since tissue disorganization extended ventrally where subretinal space was missing, we analyzed cell shape by measuring length and width of 21-25 cells per embryo directly adjacent to the subretinal space and RPE patterning defects of the ventral optic cup (temporal side, proximal level; see Supplementary Figure 2K for examples). In controls, progenitor cells are pseudostratified/columnar in both presumptive RPE and retina, with an average length/width ratio between 2.5-3.3 per embryo (SEM +/-0.16-0.26; n=3 embryos). In *Cdc42*^*CKO*^, the average of the length/width ratio is smaller, between 1.8 and 2.1 (SEM +/-0.11-0.15; n=4 embryos), with a significant difference compared to controls (Supplementary Figure 2L; p=0.0097). Thus, in *Cdc42*^*CKO*^ optic cups, disorganized regions adjacent to defective subretinal space and RPE patterning contain fewer columnar progenitor cells and more with squamous shape. To investigate a potential defect in cellular alignment, we measured the angle between the length of each cell and the basal boundary of the optic cup (set as 0 degrees; see Supplementary Figure 2K for an example). Controls showed an average angle between 72.68-78.39 degrees (SEM +/-1.34-3.03; n=3 embryos). The average angle in *Cdc42*^*CKO*^ was measured between 20.27 and 40.27 degrees (SEM +/-3.71-5.06; n=4 embryos), with a significant difference (p=0.0007; Supplementary Figure 2M). Therefore, cell shape and cell alignment are correlated with defects in subretinal space formation and abnormal RPE patterning. Our data shows that Cdc42 is essential for expression of polarity, cell adhesion and actomyosin proteins specifically during morphogenesis and coordinated tissue organization of the ventral optic cup that needs to undergo extensive growth, invaginate, and bending to facilitate approaching of the optic fissure margins.

### Proliferation and apical localization of G2/M cells are abnormal in the *Cdc42*^*CKO*^ optic cup

To examine whether loss of *Cdc42* affected earlier eye development, we analyzed E10.5 and E11.5 embryos. At E11.5, *Cdc42*^*CKO*^ eyes exhibited a severe loss of ventral optic cup tissue, resulting in a wide coloboma and microphthalmia with a significant reduction of the eye circumference by 24% (Figure 3B, C; see Methods for details on quantification of eye circumference; 13-14 eyes from 7 embryos per genotype). Consistent with E12.5, the proximal ventral RPE layer is abnormal showing RPE patterning defects and loss of subretinal space as shown by PKCZ labeling (Figure 3E, G). To determine when exactly loss of *Cdc42* starts to affect eye development, we analyzed E10.5 embryos. At this age, *Cdc42*^*CKO*^ embryos did not show obvious eye abnormalities (Figure 3H-K). However, coronal views of cryostat sections revealed that TJP1 and PARD6 expression did not fully extend into the ventrodistal optic cup indicating disrupted establishment of apical boundaries (Figure 3M, O). Thus, at E10.5 loss of *Cdc42* started to affect formation of subretinal space.

**Figure 3:**
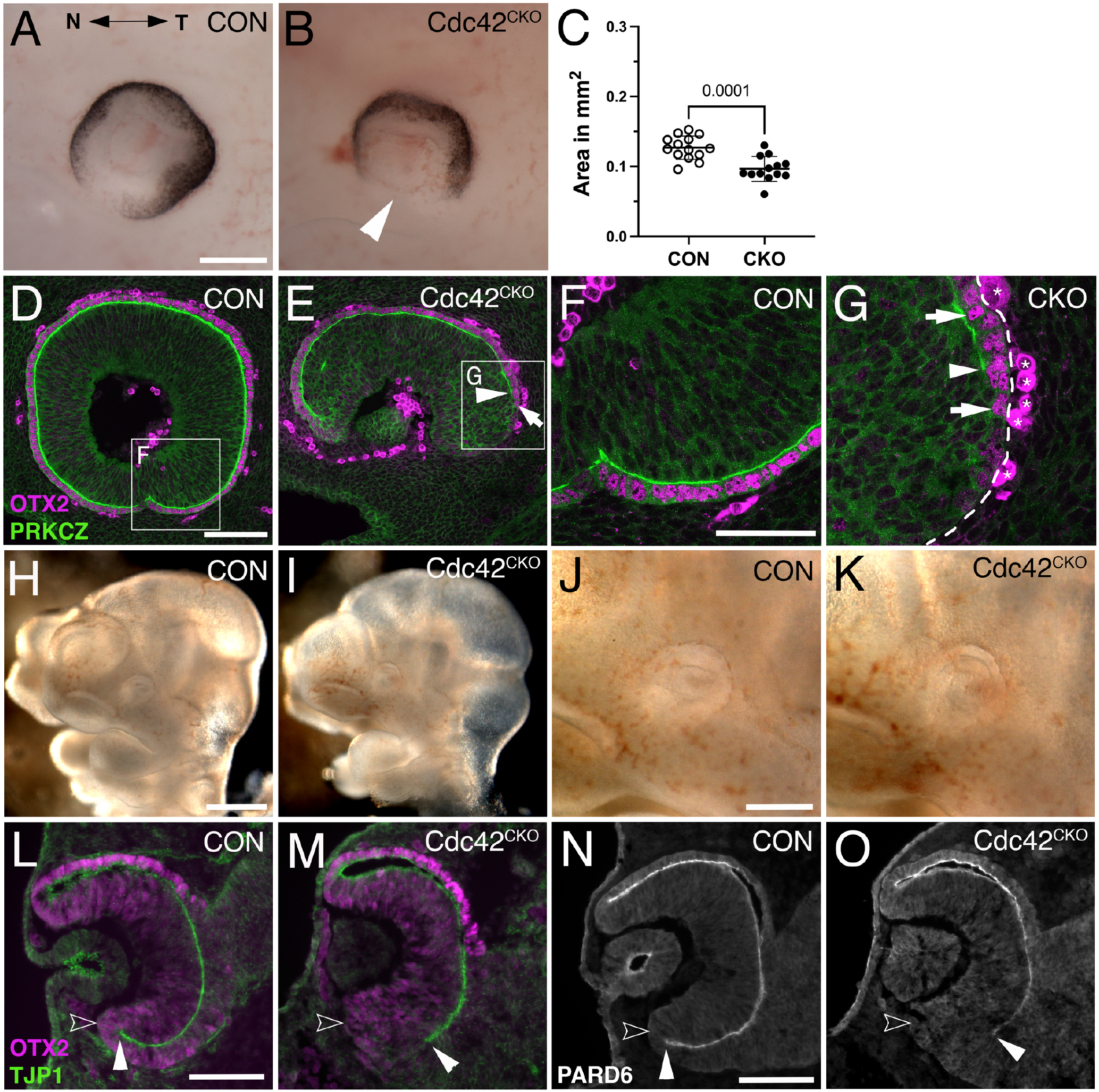
*Cdc42*^*CKO*^ optic cups are microphthalmic at E11.5, and one day earlier defects in formation of the ventral apical boundaries encompassing the subretinal space are detectable. (A, B) E11.5 optic cups. Control (A, *Cdc42*^*FL/FL*^). *Cdc42*^*CKO*^ optic cup showing a wide coloboma (B, arrowhead). C) Quantification of the circumferential eye area of left and right eyes at E11.5 shows a significant decrease by 24% in *Cdc42*^*CKO*^ (data points represent 13-14 eyes from 7 embryos per genotype, unpaired T-test). (D-G) Sagittal orientation, nasal and temporal orientation are left and right, respectively. High magnification of insets from (D, E) are shown in (F, G). Colocalization of OTX2 (magenta, arrows in E, G) and PRKCZ (green) in the temporal ventral optic cup. In *Cdc42*^*CKO*^, the apical space is shortened and does not extend into the ventral optic cup (E, G, arrowheads). Non-specific labeling on the basal side are labeled by asterisks (G). (H-K) Lateral view of E10.5 control (*Cdc42*^*FL/FL*^, H, J) and *Cdc42*^*CKO*^ embryos (I, K). *Cdc42*^*CKO*^ embryonic heads and eyes appear similar to controls. (L-O) Coronal orientation. (L-M) Co-labeling for OTX2 (magenta) and TJP1 (green) reveal apical localization of TJP1 extending very close to the distal edge of the ventral optic cup in control (L, filled arrowhead). Open arrowhead marks the outer, basal side of the optic cup. (M) In *Cdc42*^*CKO*^ embryos, TJP1 expression does not reach the distal edge in the ventral optic cup. (N, O) Localization of PARD6 is missing in the ventral *Cdc42*^*CKO*^ optic cup indicating shortening of the subretinal space (filled arrowheads). Open arrowhead marks the outer, basal side of the optic cup (N, O). Scale bars A, J: 200 µm, D, L, N: 100 µm, F: 50 µm.

To determine the underlying defect resulting in loss of ventral optic cup tissue in *Cdc42*^*CKO*^, we analyzed cell death and proliferation at E10.5. Programmed cell death occurs normally during optic cup morphogenesis (Laemle et al., 1999; Ozeki et al., 2000). The number of TUNEL-labeled cells in *Cdc42*^*CKO*^ optic cup showed no significant difference with a slight trend toward being increased (Figure 4A; Supplementary Figure S3A, B). The total number of EdU-labeled cells in the entire optic cup was significantly reduced by 10-12% in *Cdc42*^*CKO*^ (Figure 4B-D). Particularly, we observed a trend in reduced proliferation in retina, RPE, and ventral optic cup of *Cdc42*^*CKO*^ (Supplementary Figure S3C, D, F-H), compared to dorsal regions (Supplementary Figure S3E; not shown). CDC42 is required for apical localization of mitotic cells (Cappello et al., 2006; Chen et al., 2006). Consistent with this, in control embryos, mitotic retinal progenitors co-labeled for PH3.1 and PJNK are localized apically (Figure 4E; Supplementary Figure S3I-L)(Oktay et al., 2008; Ribas et al., 2012). In *Cdc42*^*CKO*^, PH3.1-/PJNK-labeled RPCs were significantly mis-localized in deeper, more basal retina regions between E10.5 and E12.5 (Figure 3F, G; Supplementary Figure S3M-N), consistent with loss of radial pseudostratified organization of retinal progenitors.

**Figure 4:**
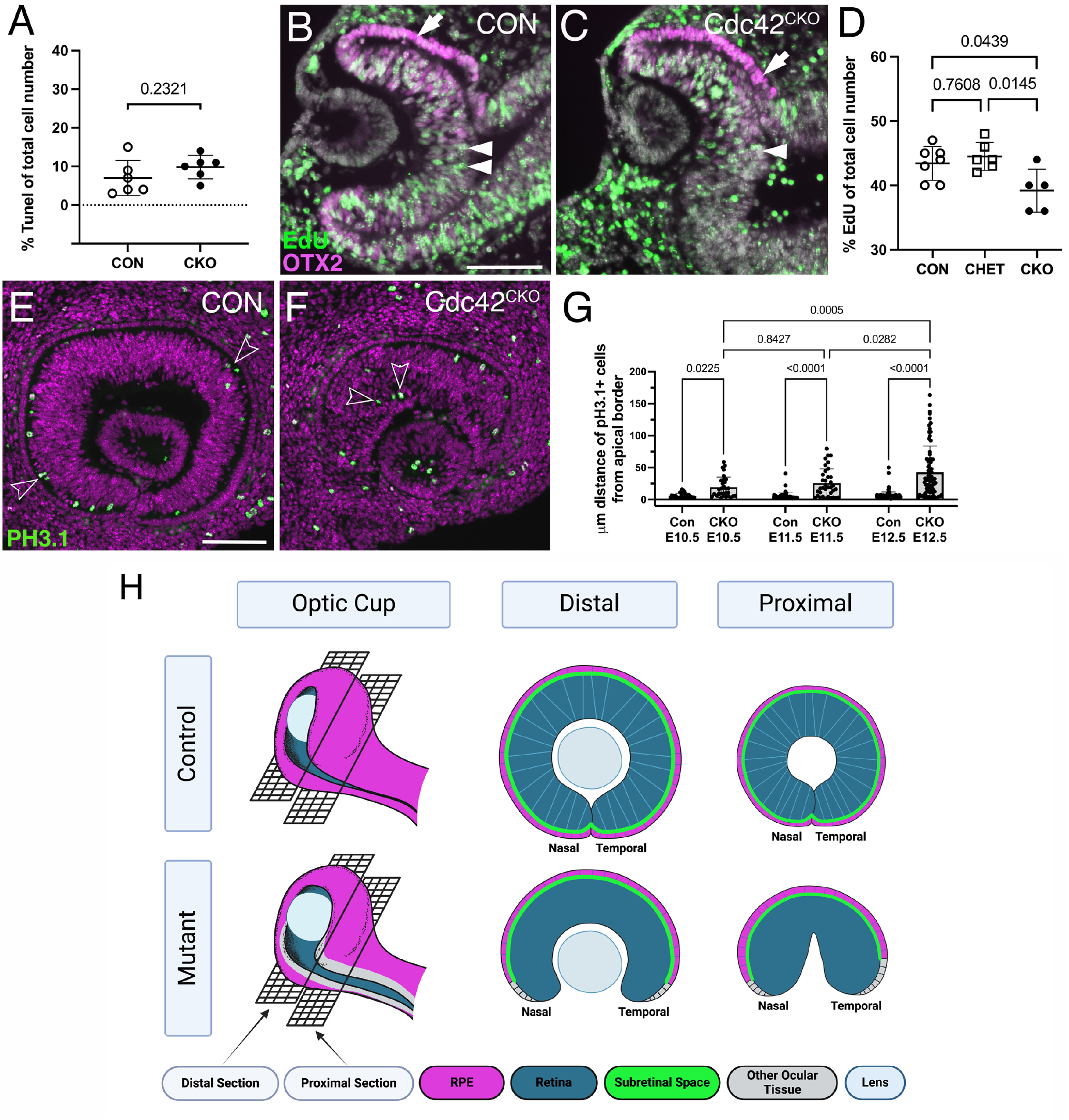
*Cdc42*^*CKO*^ eyes show defective proliferation and mis-localization of retinal progenitors. (A) Quantification of TUNEL-labeled cells shows no significant difference with a slight trend toward an increased number of apoptotic cells in E10.5 *Cdc42*^*CKO*^ optic cups. Each data point represents n=1 embryo, total of n=6 embryos for *Cdc42*^*FL/FL*^ control or *Cdc42*^*CKO*^ (unpaired T-test). (B, C) EdU incorporation in control (B, *Cdc42*^*FL/FL*^) and *Cdc42*^*CKO*^ optic cup at E10.5 (C, green, arrows). Co-labeling with OTX2 (magenta, arrow) and Dapi (grey). (D) Quantification reveals that significantly fewer cells incorporate EdU in the entire optic cup of E10.5 *Cdc42*^*CKO*^ embryos (data points plotted are number of Con n=7, CHET n=6, *Cdc42*^*CKO*^ n=5 embryos, one-way ANOVA with Tukey’s posthoc analysis). (E-G) Effect of loss of *Cdc42* on localization of PH3.1-labeled retinal progenitors in E11.5 embryos (PH3.1: green; Dapi: magenta). Normally, PH3.1-labeled cells localize at the apical boundary of the retina (E; open arrowheads). In *Cdc42*^*CKO*^ embryos, retinal PH3.1-labeled progenitors are mis-localized into deeper, more basal regions in the retina (F; open arrowheads). G) Quantification of the distance of PH3.1-labeled cells from the apical boundary reveals that mis-localization in *Cdc42*^*CKO*^ is significant, starting at E10.5 (n=3-5 embryos per genotype, 2-way ANOVA with Tukey posthoc analysis). Plotted are individual cells for each timepoint and genotype. (H) Model summarizing the observed defects with emphasis on regional domains in the optic cup. (Created in Biorender.com) For explanation, see text. Scale bars B, E: 100 µm.

## Discussion

Our results show that temporally controlled, Cre-mediated ablation of *Cdc42* at the onset of optic vesicle evagination (E8.5) caused severe morphogenesis defects of the ventral optic cup. Mitotic retinal progenitors in *Cdc42* mutants become progressively mis-localized basally and lose the pseudostratified organization of the retinal epithelium. Combined with increased apoptotic cell death and reduced proliferation, these abnormalities likely cause growth arrest of the ventral optic cup. Consequently, *Cdc42* mutant optic cups are microphthalmic and exhibit a large ventral coloboma. The ocular growth defect becomes progressively worse (reduction of eye size by 24% and 35% at E11.5 and E12.5, respectively) and is, therefore, not due to a developmental delay. Distribution of β-catenin in *Cdc42*^*CKO*^ optic cups did not reveal obvious changes suggesting that intercellular junctions may not be severely disturbed, however, we cannot exclude effects on other components of the adherens junction complex. Other studies have shown that Cdc42 is necessary to regulate proliferation (Cappello et al., 2006; Chen et al., 2006; Melendez et al., 2013); for example, mis-localization of PH3.1-labeled neuroepithelial progenitors in the forebrain of *Cdc42*^*CKO*^ is correlated with hyperproliferation (Chen et al., 2006). Conversely, during development of the heart, teeth and kidney tubules, tissue-specific ablation of *Cdc42* results in decreased proliferation of cardiomyocytes, dental mesenchyme and kidney tubule formation (Elias et al., 2015; Li et al., 2017; Ma et al., 2021), Here we report that progenitors in *Cdc42*^*CKO*^ optic cups show decreased proliferation, in combination with loss of apicobasal polarity proteins. Thus, the effect of *Cdc42* disruption on developmental proliferation is dependent on the tissue context. In developing cardiomyocytes, *Cdc42* disruption results in a decrease of cyclin B1 (Li et al., 2017), therefore, cell cycle regulators could be similarly deceased in our *Cdc42*^*CKO*^ optic cups. In addition, it may be interesting to investigate whether Cdc42 participates more directly in gene expression by regulating nuclear actin dynamics during development of the ventral optic cup (Rajakyla and Vartiainen, 2014).

Additional novel defects (to our knowledge) include cellular and tissue abnormalities in the ventral optic cup; a regionally restricted loss of expression of polarity, adherens junction proteins and actomyosin lining the apical boundaries of retina and RPE as well as key regulatory proteins for RPE regionalization. These defects in *Cdc42*^*CKO*^ optic cups correlate with absence of subretinal space between the apical boundaries of retina and RPE in the ventral-most region of the optic cup and resulting in loss of a structurally distinct RPE layer adjacent to the optic fissure. Our data here reveals a novel, very specific role for Cdc42 in regulating growth, RPE regionalization, and optic fissure closure in the ventral optic cup. The following model summarizes these defects, with emphasis on regional domains in the optic cup (Figure 4H). In *Cdc42* mutant eyes, subretinal space is absent in the ventral optic cup, adjacent to the optic fissure, and cells that should be continuous with the dorsally located RPE layer (magenta) fail to differentiate into RPE (grey). This defect is worse in the proximal optic cup, specifically in the temporal domain. The growth defects result in smaller eyes (microphthalmia) and failed closure of the optic fissure. In addition, retinal progenitors fail to maintain a pseudostratified organization.

### Disruption of *Cdc42* at the optic vesicle stage causes disorganization of the retinal neuroepithelium at E11.5, earlier than previously reported

Our study is the first to examine in detail how disruption of *Cdc42* in ocular neuroepithelial tissues affects optic vesicle and optic cup morphogenesis. Other loss of function studies revealed few roles for Cdc42 in developing ocular tissues. In mouse, it is necessary for lens pit invagination during optic cup morphogenesis by inducing filopodia to couple lens vesicle and retina (Chauhan et al., 2009). In addition, lens placodal cells going through planar polarization during invagination require Cdc42 to regulate junctional elongation (Muccioli et al., 2016). These loss of function studies were performed with lens placode-specific inactivation of *Cdc42*. Here, we used *Hes1*^*CreERT2*^ that also induces mosaic recombination in the lens vesicle, thus, we cannot exclude that filopodia formation or cell shape changes in lens placodal cells is disturbed in *Cdc42*^*CKO*^.

Furthermore, Cdc42 is required for lamination and tissue integrity during retinal neurogenesis and for maintenance of photoreceptor outer segments in mouse and zebrafish (Choi et al., 2013; Heynen et al., 2013). Particularly, loss of Cdc42 in the developing mouse retina at E10.5 resulted in severely disrupted lamination after E14.5, retinal degeneration and affected visual function postnatally (Heynen et al., 2013). In this context, it is suggested that Cdc42 is critical for formation of assembly of adherens junctions. Loss of function analyses for polarity and adherens junction proteins are consistent with a critical role in progenitor alignment and lamination in the developing retina during neurogenesis (Chan et al., 2020; Chen et al., 2013; Erdmann et al., 2003; Fu et al., 2006; Horne-Badovinac et al., 2001; Wei et al., 2004; Westenskow et al., 2009; Yamaguchi et al., 2010; Zhang et al., 2009). Our data shows that loss of *Cdc42* at E8.5 interferes with pseudostratified organization of retinal progenitors that becomes obvious at E11.5. The mis-localization of PH3.1 labeled cells in *Cdc42*^*CKO*^ suggests that abnormalities in retinal progenitor arrangement are detectable at E10.5. Our observations are consistent with and extend previous studies; we show earlier effects of loss of Cdc42 on progenitor organization in the retinal epithelium in mouse.

### Cdc42 is specifically required in the ventral optic cup but dispensable dorsally

The formation of adherens junction and apicobasal polarity of the dorsal apical boundaries of retina and RPE appears not affected in *Cdc42*^*CKO*^ optic cups. Compared to controls, we observed no obvious difference in dorsal localization of PARD6, PKCZ, F-actin, TJP1, β-catenin or pMLC2. This is unusual and suggests that Cdc42 is not required for apicobasal polarity and adherens junction formation in the dorsal optic cup.

A direct target of CDC42 is the kinase MRCK that phosphorylates pMLC2 to promote apical constriction (Zihni et al., 2017). During evagination of the optic vesicle, the actomyosin regulator pMLC2 is expressed throughout the apical boundary, and it becomes downregulated in the apical boundary of the presumptive retina during optic cup invagination (Eiraku et al., 2011). This results in differential tissue rigidity aiding in the invagination process that requires unusual basal constriction of the retina (Eiraku et al., 2011). PMLC2 is absent in the edges of the ventral *Cdc42*^*CKO*^optic cup, suggesting that reduced contractibility and loss of apical tension may contribute to preventing apposition of optic cup margins in *Cdc42*^*CKO*^.

### Failure of RPE differentiation does not result in transdifferentiation into retina

The ventral *Cdc42*^*CKO*^ optic cup showed reduced MITF and OTX2 protein expression, especially in the proximal domain. Indeed, a defined, separate layer continuous with the dorsal RPE is missing in the ventral *Cdc42*^*CKO*^ optic cup and this was closely correlated with an absence of PARD6, PRKCZ, TJP1, F-actin and pMLC2. These observations suggest that presumptive retina and RPE cells adjacent to the ventral fissure fail to establish apicobasal polarity and do not form two separate apical boundaries encompassing subretinal space. Normally, the subretinal space may be necessary for cells in the presumptive RPE layer to be separated from retina-inducing signals (e.g. FGF) to properly differentiate. However, in *Cdc42*^*CKO*^ signals from the extraocular mesenchyme may not be sufficient to promote RPE differentiation. In many genetic mouse mutants showing defective RPE fate during optic cup morphogenesis, the tissue adopts neural retina fate via transdifferentiation (reviewed in (Fuhrmann et al., 2014). In *Cdc42* mutant optic cup, retina-specific Vsx2 expression is not expanding into the area of RPE loss suggesting that RPE-to-retina transdifferentiation does not occur. We cannot exclude that additional time may be needed for transdifferentiation to occur in older embryos. It is also likely that failure of RPE differentiation in the ventral optic cup margins by itself may interfere with optic fissure closure, as shown in other studies (Boobalan et al., 2022; Cai et al., 2010).

### Signaling pathways and Cdc42 regulation of ventral optic cup morphogenesis

Altogether, our data reveals a severe complex phenotype of defective eye morphogenesis, caused by *Cdc42* disruption. Particularly, growth and optic fissure closure of the ventral optic cup is prominently affected, similar to genetic manipulation of Wnt, Hippo, FGF, BMP, hedgehog pathways or transcription factors such as Pax2, Vax1/2, among others (Alldredge and Fuhrmann, 2016; Bankhead et al., 2015; Boobalan et al., 2022; Cai et al., 2013; Chen et al., 2013; Fuhrmann et al., 2022; Hocking et al., 2018; Knickmeyer et al., 2018; Lahrouchi et al., 2019; Mui et al., 2005; Sun et al., 2020; Yan et al., 2020; Zhou et al., 2008)(for review, see (Peters, 2002)). These factors may affect mostly the ventral optic cup domain since it undergoes more complex and dramatic changes in morphology at this time. Specifically, the ventral domain needs to undergo extensive growth and bending to facilitate approaching of the optic fissure margins. In other developmental systems, Cdc42 has been shown to execute diverse cellular processes downstream of signaling pathways, for example, Wnt/Ror2 or Hippo/Yap (Hikasa et al., 2002; Ma et al., 2021; Sakabe et al., 2017). Thus, Cdc42 represents an important effector downstream of identified key regulators during optic cup morphogenesis. Furthermore, we observed overall a differential effect of *Cdc42* loss on nasal and temporal domains in the ventral optic cup.

Differential differentiation of nasal and temporal domains is important during the development for high acuity vision. Differential growth of nasal and temporal domains in the optic cup is regulated by signaling pathways upstream of domain-specific transcription factors, as shown in zebrafish, chick, and mouse (Hernandez-Bejarano et al., 2022; Hernandez-Bejarano et al., 2015; Schulte and Cepko, 2000; Smith et al., 2017). We reported recently that deletion of neurofibromin 2 results in thickening and hyperproliferation of the ventro-temporal RPE layer (Sun et al., 2020). Thus, the temporal domain in the ventral optic cup could be also more sensitive to disturbances during development.

In humans, a failure of the optic fissure to close occurs between 5-7 weeks of gestation, with an incidence of up to 10% of childhood blindness (Patel and Sowden, 2019). Depending on where and when coloboma manifests, it can result in impairment of visual function. The optic cup morphogenesis defects that we observed in *Cdc42*^*CKO*^ results in a phenotype resembling chorioretinal coloboma, missing retina, RPE and choroid. Coloboma in humans can be accompanied with microphthalmia, and the *Cdc42*^*CKO*^ is microphthalmic. While many genes causative for coloboma have been identified, in most cases the genetic cause is unknown. Thus, Cdc42 may be an important gene regulating optic fissure closure in humans.

## Acknowledgements

We are grateful to members of the Fuhrmann laboratory for their technical support and helpful comments. We appreciate advice and constructive discussions from members of the Levine laboratory. We thank Charles Murtaugh (University of Utah, Salt Lake City, UT) for kindly providing the mouse line *Hes1*^*CreERT2*^.

