## Supplementary figures and images for "Loss of Cdc42 causes abnormal optic cup morphogenesis and microphthalmia in mouse"

### Supplemental Figure 1

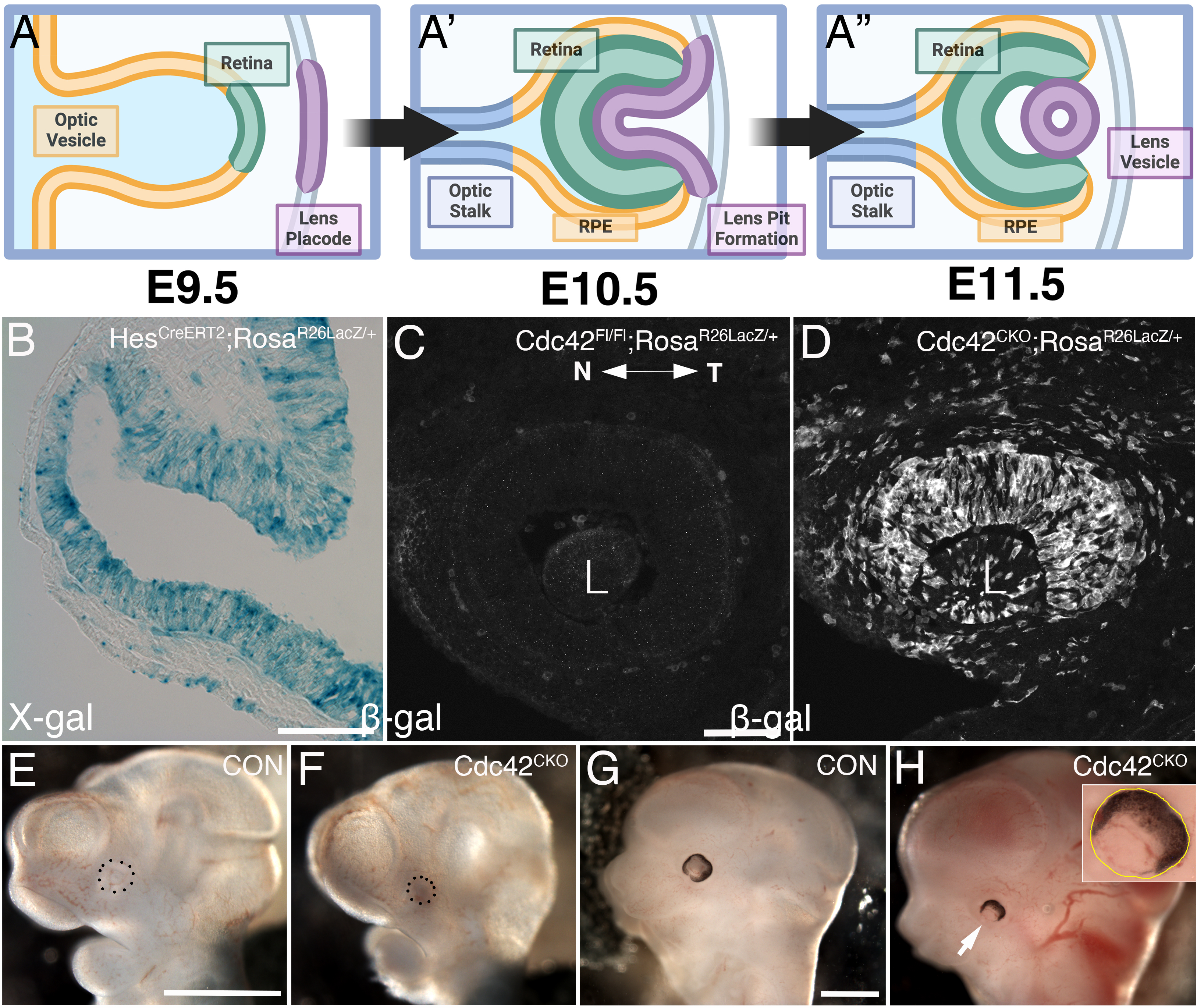

### Supplemental Figure 2

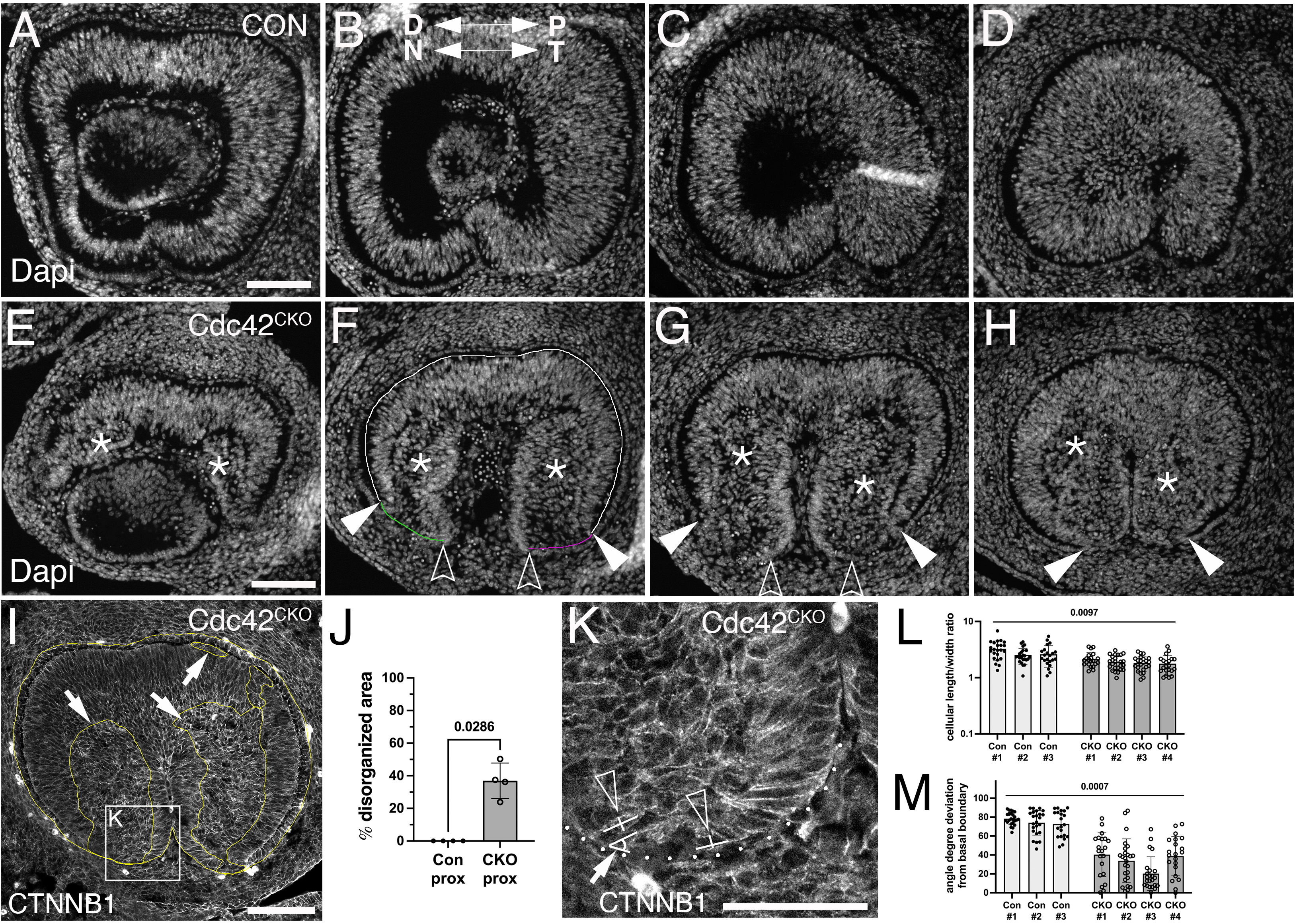

### Supplemental Figure 3

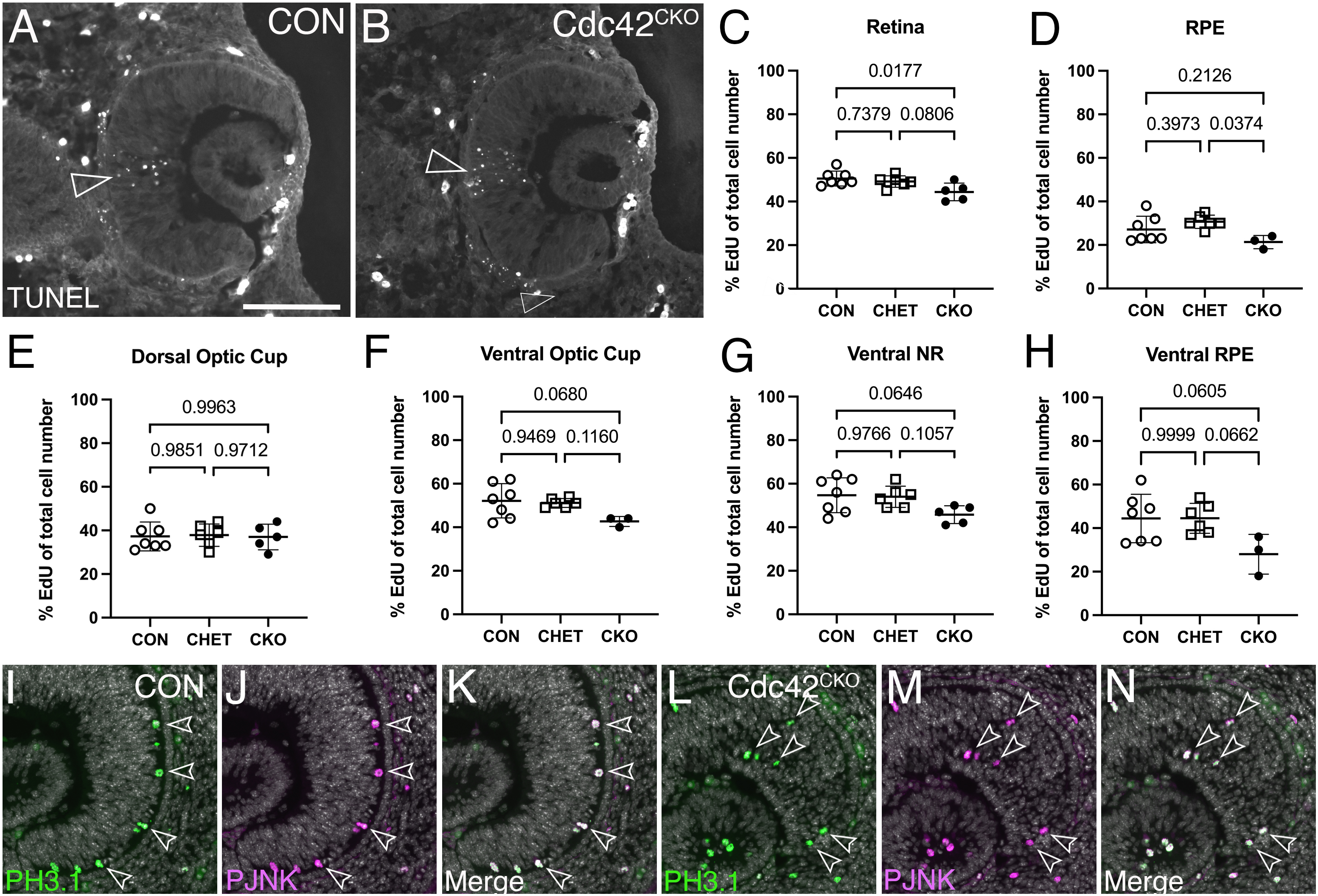
